# Single-molecule mapping of replisome progression

**DOI:** 10.1101/2022.01.10.475700

**Authors:** Clémence Claussin, Jacob Vazquez, Iestyn Whitehouse

## Abstract

Fundamental aspects of DNA replication, such as the anatomy of replication stall sites, how replisomes are influenced by gene transcription and whether the progression of sister replisomes is coordinated are poorly understood. Available techniques do not allow the precise mapping of the positions of individual replisomes on chromatin. We have developed a new method called Replicon-seq that entails the excision of full-length replicons by controlled nuclease cleavage at replication forks. Replicons are sequenced using Nanopore, which provides a single molecule readout of long DNA molecules. Using Replicon-seq, we have investigated replisome movement along chromatin. We found that sister replisomes progress with remarkable consistency from the origin of replication but function autonomously. Replication forks that encounter obstacles pause for a short duration but rapidly resume synthesis. The helicase Rrm3 plays a critical role both in mitigating the effect of protein barriers and facilitating efficient termination. Replicon-seq provides an unprecedented means of defining replisome movement across the genome.

## Introduction

In eukaryotes, DNA replication originates from multiple sites along chromosomes known as origins of replication. The replicative helicase – CMG – is coupled to several other replication factors and the replicative DNA polymerases to form the replisome, which collectively synthesizes the nascent genome (Burgers and Kunkel, 2017). Following initiation, two sister replisomes emanate from a replication origin and progress along the DNA template in opposite directions until they meet a DNA end, or a convergent replisome where a “termination” process results in CMG unloading and the ligation of two replicons (Dewar and Walter, 2017). During S-phase, sister replisomes can remain in proximity (Kitamura et al., 2006) and may be physically coupled (Yuan et al., 2019) which could allow some degree of coordination of their movement (Conti et al., 2007; Li et al., 2020; Yuan et al., 2019). However, *in vitro* evidence indicates that replisomes can function as monomers (Yardimci et al., 2010). Whether sister replisomes are functionally coupled *in vivo* – that is: a stall of one replisome is communicated to its sister – is poorly understood and has not been definitively addressed.

The study of DNA replication at nucleotide resolution with genomics has significantly lagged behind the transcription/RNA fields. This is principally because nascent DNA molecules are limited in copy number, differ in length by several orders of magnitude and are difficult to unambiguously separate from their template. Population based sequencing methods to study DNA replication are broadly insensitive to coincident or stochastic events affecting the replisome. Recent advances such as single-cell replication profiling (Chen et al., 2017; Dileep and Gilbert, 2018), high throughput combing (Wang et al., 2021), and nucleotide analogue detection with nanopore sequencing (Hennion et al., 2020; Muller et al., 2019) have begun to reveal DNA replication patterns at single molecule resolution. Yet, available methods have limited utility in precise mapping of replisome locations and are unsuitable for the fine scale analysis of replication defects.

We describe a new method which is able to overcome the many of the limitations of genome-wide approaches to study DNA replication. We utilize long-read Nanopore sequencing to identify and sequence nascent DNA molecules in their entirety. This approach maps the ends of nascent DNA molecules with nucleotide resolution and allows the precise localization of replisomes along the genome. We show that sister replisomes progress through tens of kilobases of chromatin at remarkably similar rates, we also find that a stall in one replisome is not communicated to its sister – indicating that replisomes function autonomously. Finally, we demonstrate that the DNA helicase Rrm3 performs a critical role in facilitating replication termination.

## Results

### Replicon-seq to study sister replication fork movement by sequencing full length replicons

We aimed to extract and sequence the DNA that has been synthesized by two replisomes that originated from the same origin of replication (replicon). We reasoned that creation of DNA strand breaks at functional replisomes would allow the position of any replisome to be identified by mapping the ends of nascent DNA by sequencing. Moreover, the relative positions of sister replisomes could be identified if nascent DNA chains could be sequenced in their entirety (Figure 1A). Given that fusion of Micrococcal Nuclease (MNase) to candidate proteins allows the generation of specific DNA breaks (Schmid et al., 2004) including at replication origins (Foss et al., 2019), we chose to fuse MNase to the Mcm4 subunit of the replicative helicase, CMG, which encircles the leading strand template and is likely the most stable component of the replisome. Consistent with reports using MNase fusions (Schmid et al., 2004; Skene and Henikoff, 2017; Zentner et al., 2015), we found no growth defect of strains cultured at 25°C and 30°C in the absence of calcium (Figure S1). We tested whether the MNase fusion had the desired specificity by inducing DNA cleavage at origins in G1 (Figure S1B); surprisingly, we found that the C-terminal fusion give rise to far more DNA cleavage than the N-terminal fusion (Figure S1B). This difference likely reflects the N-N loading of two MCM2-7 hexamers at origins (Douglas et al., 2018; Georgescu et al., 2017) which may restrict access to the DNA of the N-terminal MNase fusion. Given that after helicase activation, the N-termini of a single MCM2-7 hexamer will be at the front of the replisome and no longer complexed with a second hexamer (Douglas et al., 2018; Georgescu et al., 2017), we reasoned that the use of the N-terminal fusion should provide a means of promoting parental DNA cleavage by MNase at “activated” helicases, rather than those that are loaded, but inactive.

**Figure 1:**
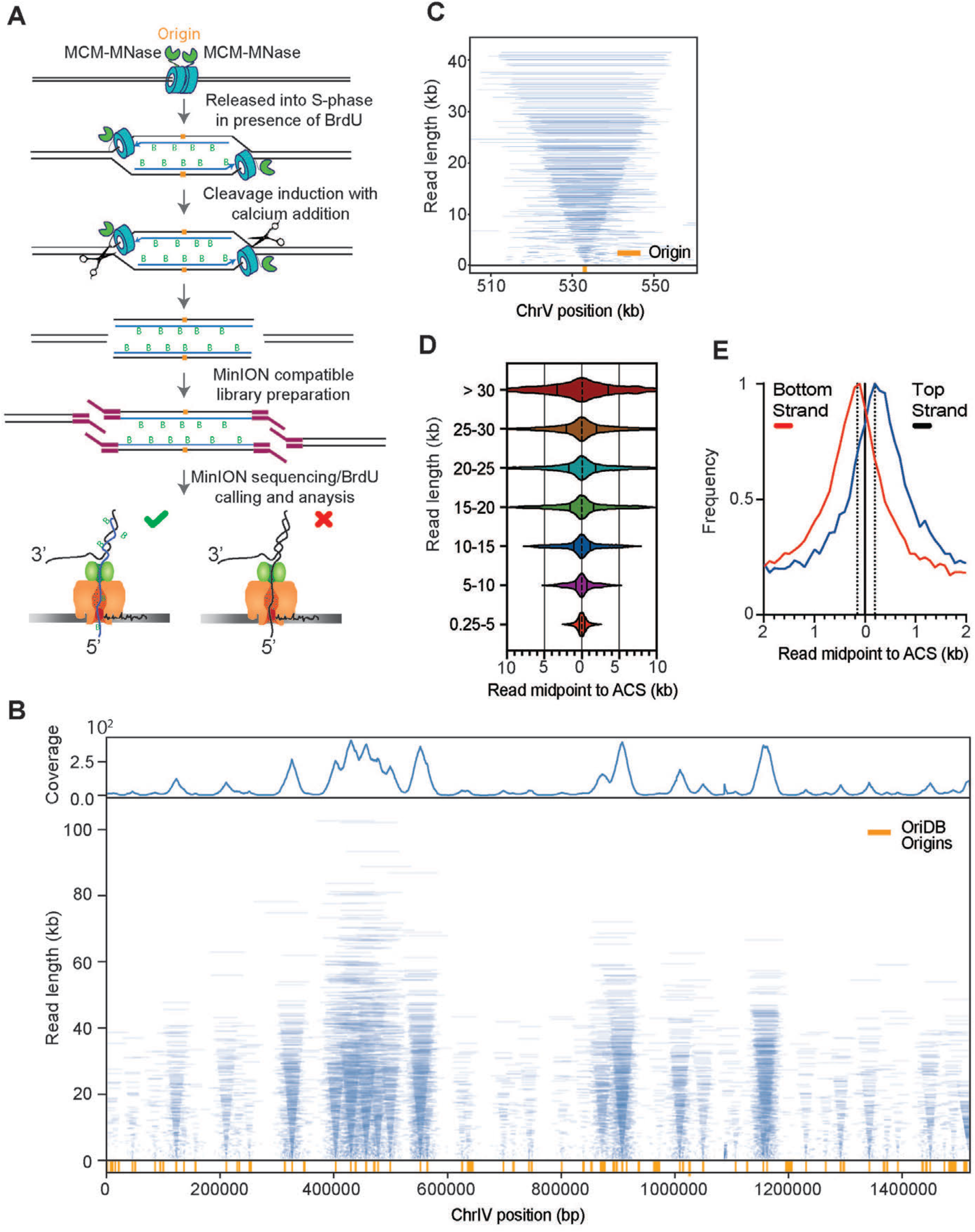
Replicon-seq a new method to study sister replication for movement. **A**. Schematic of Replicon-seq methodology. **B**. Tornado plot of Replicon-seq data. BrdU containing reads are shown, each line represents a single read graphed by length and chromosome position, line transparency is scaled to reflect read abundance. Genome coverage of the displayed reads is shown on the upper track. Replication origins defined by OriDB are shown on the lower track. **C**. Zoom of representative origin (ARS520) showing symmetry of replicons, each blue line represents an individual read. **D.** Violin plot showing the positions of read midpoints relative to the ACS at replication origins for different length reads. **E**. Anchor plot showing the position of the midpoint of Top and Bottom-strand reads centered on the ACS.

We performed experiments in which *S. cerevisae* are arrested in G1 with alpha-factor and BrdU is added to allow labeling of nascent DNA upon release into S-phase. Cells were harvested in early S-phase, permeabilized and MNase activated with the addition of calcium for limited time, on ice (Methods). The DNA is directly sequenced using nanopore technology (Lu et al., 2016) with the output expected to be composed of nascent – BrdU containing molecules – as well as parental template and un-replicated DNA. BrdU containing reads were selected informatically (Boemo, 2021; Muller et al., 2019) (Methods).

Nanopore sequencing of replicons proceeds from the 5’ end of the lagging strand through to the 3’ end of the leading strand of the sister replisome (Figure 1A). Thus, the ends of replicons should provide a high-resolution, single-molecule readout of the relative position of sister replisomes along the genome. Plotting individual BrdU-containing reads according to genomic location and length reveals highly symmetric patterns which emanate from known replication origins (“tornado” Figure 1B; Figures S2, S3). The median read-length of BrdU containing molecules is ∼10kb but many molecules are greater than 50kb (Figure S1C). Small molecules are also present, and typically cluster around replication origins (Figure 1B, Figure S2, S3), the genesis of this DNA is unclear, but most likely reflect sequencing of broken, or nicked replicons.

The symmetric pattern of replicons shown in Figure 1, B and C should only be apparent if two sister replisomes move at a similar rate. Indeed, provided that replisome progression is consistent, the midpoint of each replicon should overlap the origin from which it was derived. Using this logic, we can map replication origins (Figure S1D) and define the consistency of sister-replisome progression by comparing the distances each sister replisome traveled from their origin (Figure 1D) (Brewer and Fangman, 1987; Eaton et al., 2010; Nieduszynski et al., 2007). Although such measurements are potentially confounded by the variability in replication initiation sites, sequencing of truncated replicons, and replicon asymmetry brought about by termination; we find that the majority of sister replisomes progress at similar rates, regardless of distance traveled: the median deviation is ∼1% and the distance traveled by 50% of all replisomes is within ∼15% of their sister (Figure 1D). The remarkable consistency in replisome rate in WT cells does not prove that progression of sister replisomes is coordinated (Conti et al., 2007; Li et al., 2020), but does indicate that robust mechanisms exist to ensure consistent replisome progression through tens of kilobases of chromatin (Bellush and Whitehouse, 2017). Through further analysis, we also find that the synthesis of the contiguous lagging strand is typically delayed by ∼350nt as compared to the leading strand (Figure 1E), a finding consistent with EM imaging (Lopes et al., 2006) and the expected size of un-ligated Okazaki fragments (Smith and Whitehouse, 2012). Our ability to detect a difference in position of leading and lagging strand ends indicates that MNase’s distinct preference for cleavage of single stranded DNA (Tucker et al., 1978), would likely result in preferential cleavage of the parental template - which is single-stranded at the replication fork. Such a cleavage pattern would likely occur under our tightly controlled reaction conditions (0°C, 10 seconds) and would preserve the integrity of the nascent DNA chains. Importantly, provided that the lagging strand template is cut, the 5’ ends of nascent DNA would be made accessible for sequencing which would allow us to assay both ends of nascent molecules (Figure S4).

### Fob1-mediated replication fork pausing at rDNA

To test if Replicon-seq can identify sites of replication fork pausing, we analyzed the well-characterized (Replication Fork Block) RFB pausing site within the rDNA locus. The presence of Fob1 bound to the DNA results in a unidirectional pausing of replisomes to prevent convergence of RNA polymerase I and the replisome in rDNA repeats (Figure 2A) (Brewer and Fangman, 1988; Brewer et al., 1992; Kobayashi et al., 1992). As anticipated, we detected a pronounced accumulation of replicon ends directly adjacent to the annotated RFB, whereas progression of the sister replisome is apparently unimpeded (Figure 2B). We analyzed how DNA synthesis of the leading and lagging strands can proceed to the RFB. Although Fob1 can bind three closely spaced motifs: RFB1, RFB2 and RFB3, the highest preference is for RFB1 (Kobayashi, 2003). We found that the lagging strand 5’ ends primarily accumulate ∼ 45-65nt upstream of RFB1 (Figure 2 C,E). This pattern may indicate that initiation of Okazaki fragments does not occur within ∼40nt of the leading edge of the replication fork, similar to previous measurements (Duxin et al., 2014; Vrtis et al., 2021). The 3’ ends of the leading strand show a similar pattern to the 5’ ends of the lagging strand: a pronounced enrichment ∼40-60nt upstream of RFB1, a distance which is likely defined by the footprint of CMG on the leading strand template (Figure 2 D,F) (Duxin et al., 2014). A significant fraction of the leading-strand 3’ ends extend beyond the RFB; it is noteworthy that extension through RFB becomes more prominent as replicons increase in size (Figure 2 D, F). Since rDNA repeats are replicated in clusters (Pasero et al., 2002), the majority of replisomes stalled at RFB soon encounter a converging replisome from an adjacent repeat. Thus, leading strand extension of longer replicons, through the RFB, likely occurs during termination; interestingly, bypass is rarely observed by the lagging strand, (Figure 2E) which may be indicative of the termination mechanism (Figure 5F). We also detect potential stall sites at the 3’ end of the 5S gene and ∼150nt upstream of RFB1; these sites contain long poly dA-dT stretches of DNA, which may cause replisome/polymerase stall (Hile and Eckert, 2008; Tubbs et al., 2018) or may be aberrant cleavage by MNase (Tucker et al., 1978).

**Figure 2:**
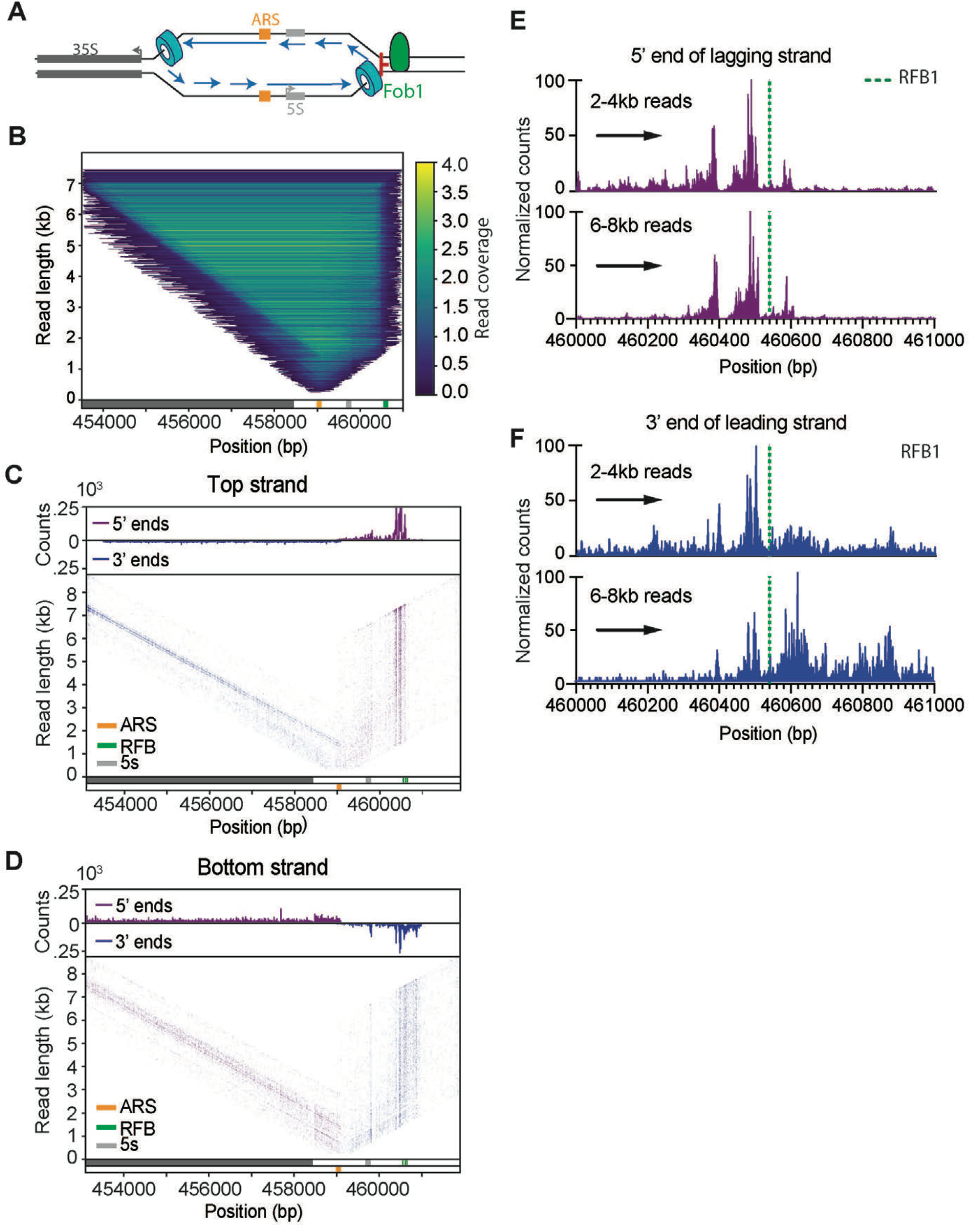
Replication fork block at rDNA locus. **A**. Schematic of rDNA locus showing the directional RFB mediated by Fob1. **B.** Heatmap of reads overlapping rDNA origin of replication (n= 5.58^^4^), each line represents an individual read, color is scaled to reflect density. **C**, **D** 5’ and 3’ DNA ends are plotted for of either Bottom or Top-strand reads at the rDNA locus. Upper tracks show the genome coverage of the 3’ or 5’ ends of reads. Prominent rDNA features are displayed in the lower track. **E**, **F** Counts of 5’ and 3'ends of the lagging or leading strands of replisomes approaching the RFB from the rightward moving fork (indicated by black arrow). Upper graphs show 2-4kb reads, lower show 6-8kb reads.

### Natural replication fork pausing sites in WT cells

Elements such as protein:DNA complexes, highly transcribed genes, R:loops and DNA structures can pose a significant challenge to the passage of the replisome and induce replisome stalling events (Shyian and Shore, 2021; Zeman and Cimprich, 2014). However, basic parameters such as the duration of the stall and the fate of the sister replisome remain poorly addressed. We therefore probed whether fork stalling in non-repetitive sequence could be detected by Replicon-seq. As shown in Figure 3, we find evidence of replisome stall in WT cells, which results in the accumulation of replicon ends at sites such as tRNA and centromeres (Deshpande and Newlon, 1996; Greenfeder and Newlon, 1992) (Figure 3A, B). Moreover, by following the progression of replicon ends past the impediment we can conclude that the replisome pauses – rather than permanently stalls at these sites. Importantly, progression of the sister replisome is not apparently affected by the stall, meaning that sister replisomes can move independently of each other, which confirms and extends previous findings (Doksani et al., 2009; Yardimci et al., 2010). We performed a meta-analysis of replicon-end accumulation across the genome to identify ∼216 stall sites in early S-phase (Methods; Figure S5A). This analysis found tRNA, centromeres, origins and 208 genes transcribed by RNA polymerase II. We performed nascent RNA-seq in early S-phase to measure transcription but found that pause sites were not generally enriched for highly transcribed genes (Figure S5B). Closer inspection showed that many pause sites in genic regions are explained due to proximity to known impediments such as centromeres and tRNA. To directly assay the effect of RNA polymerase II transcription, we focused on the PDC1 gene whose transcription is >6 standard deviations above the S-phase mean (Figure S5B), which presumably increases the likelihood that the replisome will frequently encounter the transcription machinery. As shown in Figure 3, C and D, *PDC1* shows a defect in replisome progression towards 5’ end of the body of the gene, nevertheless, even at *PDC1* replisomes experience modest slowing and quickly resume synthesis at a similar rate to the sister (Figure 3D), illustrating the efficient mechanisms employed to mitigate transcription/replication conflicts in WT cells (Gomez-Gonzalez and Aguilera, 2019).

**Figure 3:**
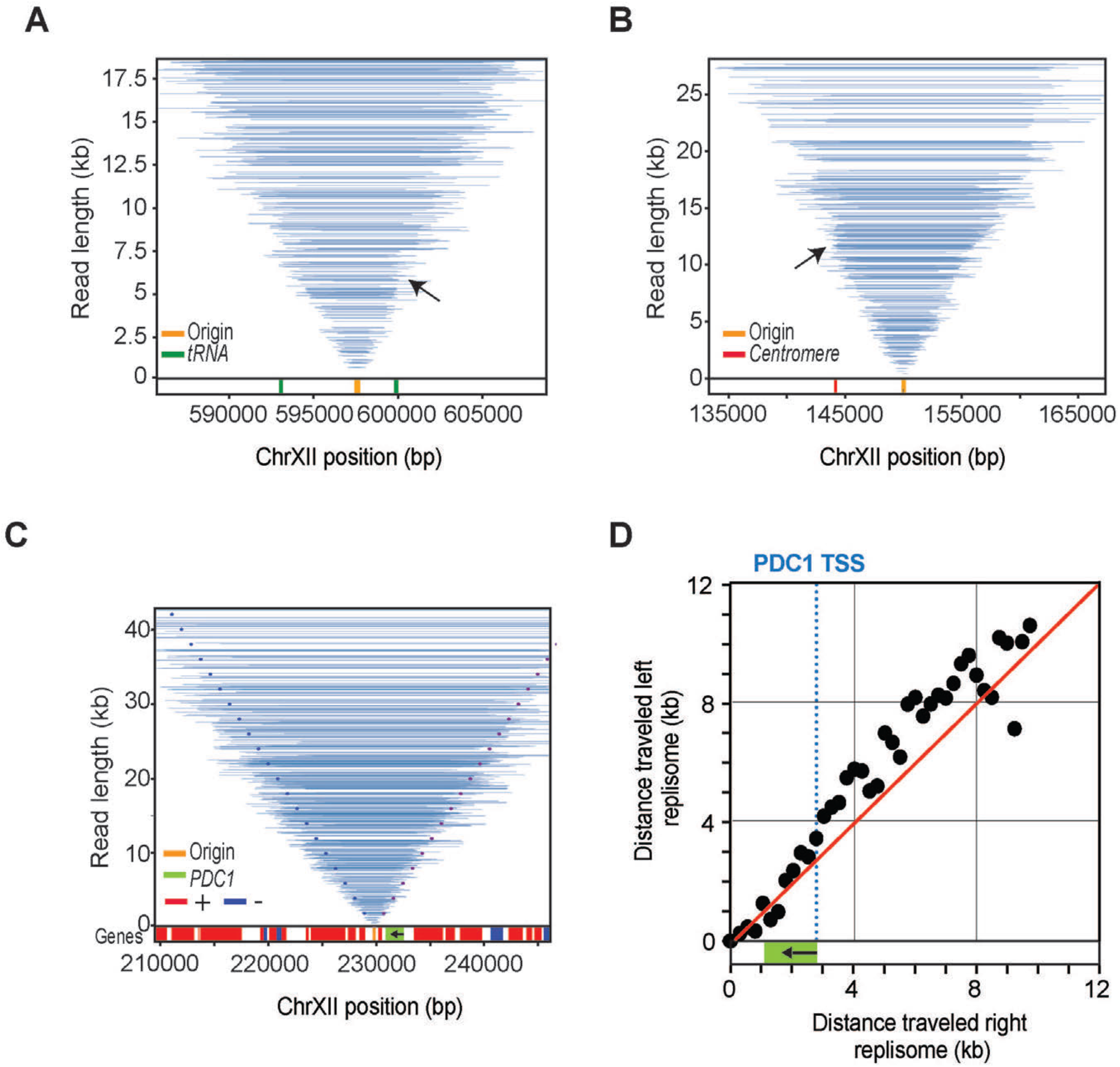
Replication fork movement in WT. Tornado plots show Replicon-seq reads and pausing events, for clarity, only reads overlapping the central origin are shown **A.** Pausing events at tRNA indicated by the black arrow, tRNA location is depicted in lower track in green. **B.** Same as A, except the arrow in B is showing the *Cen12* (lower track red). **C.** Same as A&B for pausing at *PDC1* gene. The dotted lines represent expected positions of read ends if left and right replisomes progress at equal rates (left end in dark blue, right end in purple) from the origin. Gene position is depicted in lower track. *PDC1* is green, arrow indicates transcription orientation. **D**. Median position of read ends in 250bp intervals was calculated for different length replicons emanating from the replication origin. The relative positions of the left and right replisomes are plotted at increasing distance from the origin. Dashed blue line represents the *PDC1* TSS. Red line shows the expected trajectory if both replicons are moving at the same rate.

### Rrm3 prevents replication fork pausing at t-RNAs and centromeres

The Pif1 family helicases – Rrm3 and Pif1 –promote replisome progression through a variety of elements, including protein DNA complexes, highly transcribed genes, tRNA and DNA structures such as G4 DNA (Muellner and Schmidt, 2020). We performed replicon-seq in *rrm3*Δ mutant strains and found that overall replication patterns appeared similar to WT, although with increased small or broken replicons (Figure 4A, Figure S6). Closer inspection of the tornado plot reveals evidence that replisomes stall at tRNA, centromeres and other prominent sites, consistent with earlier reports (Figure 4B, Figure S7A) (Ivessa et al., 2003). Replisome stalls in *rrm3*Δ persist longer than WT, yet similar to WT, replication resumes past the stall site and progression of the sister replisome appears unaffected (Figure 4A, B) – a result that clearly demonstrates that forks do not terminally arrest at tRNA in *rrm3*Δ. We developed a protocol to measure replicon asymmetry induced by replisome stalling (Figure 4C), allowing us to plot the extent a replisome progresses whilst its sister is paused. Figure 4D shows a comparison of progression asymmetry of WT and *rrm3*Δ at tRNA in both head-on or co-directional orientations with respect to tRNA transcription. Assuming that replisomes travel ∼2.0 kb per minute (Sekedat et al., 2010), we find that the pause duration in WT cells is typically less than 1 minute (Deshpande and Newlon, 1996), whereas the pause in *rrm3*Δ can extend past 4 minutes. We note that head-on conflicts take longer to resolve than co-directional conflicts – consistent with previous reports (Deshpande and Newlon, 1996; Ivessa et al., 2003; Osmundson et al., 2017; Tran et al., 2017). The abundance of stall events at tRNA genes in the *rrm3*Δ mutant allowed us to more closely investigate the specific replisome impediment. Figures 4E shows co-directional encounters result in a stall upstream of the tRNA with a prominent accumulation of 5’ nascent ends ∼40–130nt upstream of TFIIIB binding site (Nagarajavel et al., 2013). Anti-directional encounters result in a similar accumulation of replicon ends downstream of the TSS/TFIIIB confirming that a primary impediment for replication through tRNA is likely TFIIIB (Yeung and Smith, 2020) (Figure S7B). Besides tRNA we identified prominent replisome stall events at all 16 centromeres, replication origins/dormant replication origins and replication termination zones (Azvolinsky et al., 2009; Fachinetti et al., 2010) (Figures 4F, Figure S7). However, only a minority of stalls occurred at genes transcribed at RNA polymerase II, leading us to conclude that Rrm3 plays a comparatively minor or infrequent role in mitigating transcription/replication conflicts(Osmundson et al., 2017). Notable exceptions are very highly transcribed genes such as *PDC1* (Azvolinsky et al., 2009) where stalls also occur at adjacent dormant origins (Figure 4F). At *PDC1*, we find clear evidence of a distinct replisome stall at the TSS – indicating that Rrm3 is likely needed to help the replisome overcome the RNA polymerase II PIC.

**Figure 4.**
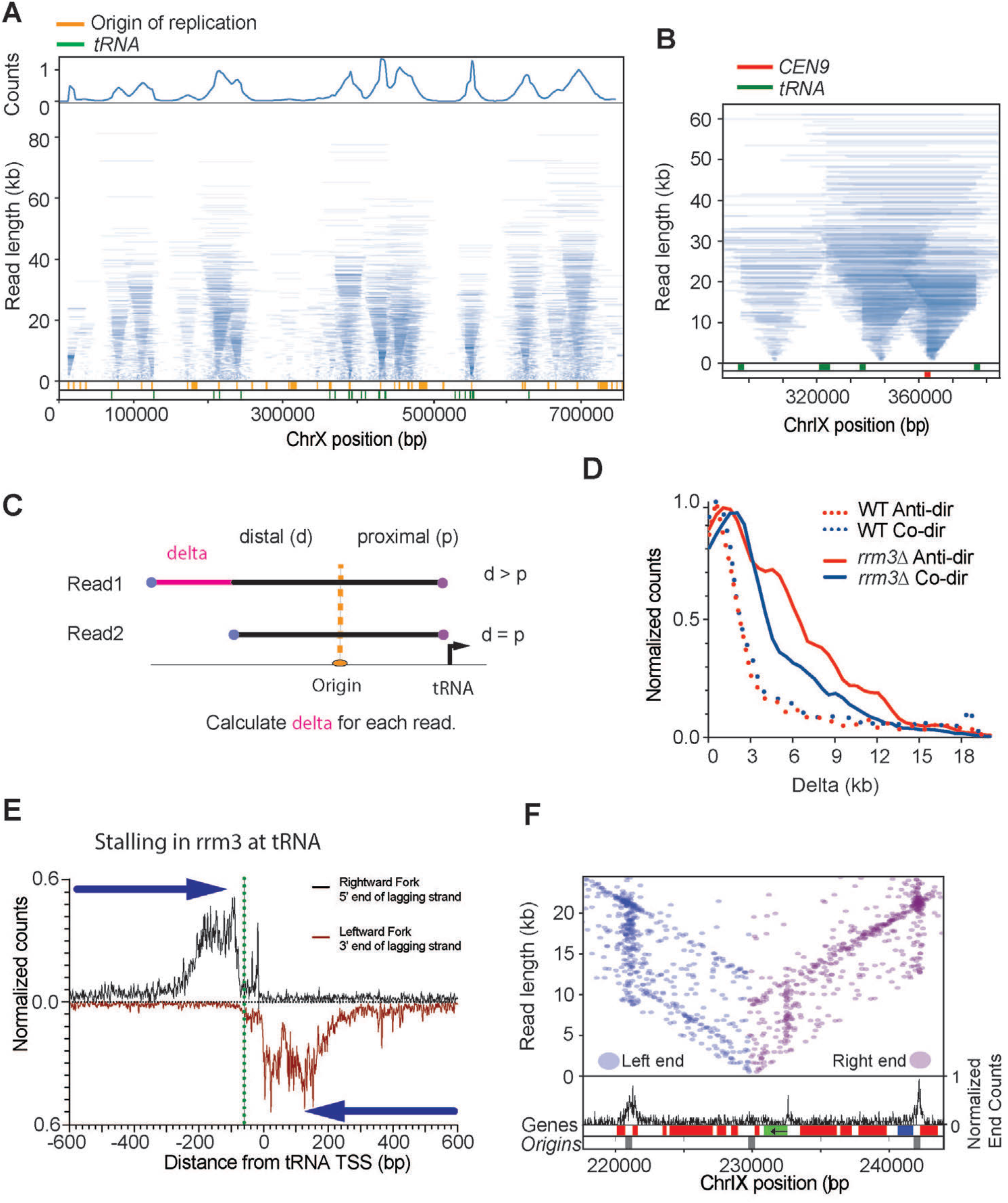
Replication fork movement in rrm3Δ **A.** Tornado plot of *rrm3*Δ strain, genome coverage is shown in the upper track, Origins of replication and tRNA locations are depicted in the lower track (orange and green respectively). **B.** Enlargement of Tornado plot to show replication fork pausing at *CEN9* (red) and tRNA (green). **C.** Schematic representing the calculation of replicon asymmetry at tRNA. **D**. Frequency plot showing extent of replicon asymmetry (delta) at tRNA genes for WT and *rrm3*Δ when tRNA are transcribed in a co-directional and anti-directional orientation with respect to the approaching replisome. **E**. Meta-plot showing the distance of the 5’ of the rightward moving fork lagging strand (black) or the leftward moving fork of the lagging strand 5’ end (red) near the transcription site of 253 tRNA genes in *rrm3*Δ. Green dotted line shows the upstream edge of TFIIIB. **F.** Read end positions for replicons overlapping ARS1211 (left ends in blue, right ends in purple). Black graph shows read-end density across the highlighted region. Gene positions (red=Bottom, blue=Top strand) and replication origins (grey) are shown at the bottom; PDC1 gene is shown in green, arrow denotes transcription orientation. Note increased read-end density i.e. stall sites, at the 5’ end of *PDC1* as well as the two origins flanking the central origin ARS211.

### Replication Termination

Replication termination zones are broadly dictated by the relative position and firing times of adjacent replication origins (Hawkins et al., 2013; McGuffee et al., 2013). In Replicon-seq, termination is evident as the fusion of two converging replication forks to produce a nascent molecule whose length is the sum of two joined replicons (Figure 5A and B). Converging replisomes do not appear to significantly slow (as compared with their sister replisomes, Figure S8) indicating that replisomes do not interfere with each other’s progress as they approach termination *in vivo*, as suggested *in vitro* (Dewar et al., 2015).

We noted that some stall sites detected in *rrm3*Δ overlap with replication termination zones (Fachinetti et al., 2010; McGuffee et al., 2013). Inspection of these sites provides evidence that adjacent replicons fail to terminate correctly in *rrm3*, resulting in an accumulation of replicon ends in the termination zone (Fig. 5C and D, Figure S6). The delay in termination in *rrm3*Δ, appears to be impacted by two phenomena: first, tRNA, centromeres and dormant origins function as bi-directional barriers to replication fork progression – meaning that termination (and joining of nascent strands) cannot occur until the barrier is passed (Figure 5C), (Fachinetti et al., 2010). Second, even in the absence of known replication barriers, converging replication forks often fail to terminate in the termination zone (Figure 5D, Figure S6), indicating that Rrm3 is required for a specific step in termination *in vivo*, consistent with *in vitro* studies and the detection of late replication intermediates on plasmids or at the rDNA (Deegan et al., 2019; Ivessa et al., 2000; Osmundson et al., 2017). The low coverage and diffuse nature of termination within the single copy genome prevented high-resolution analysis of termination; thus, we chose to investigate termination at the rDNA in *rrm3*Δ. Focusing on long replicons, which should be enriched for molecules in the process of termination, we calculated the read coverage on the leading and lagging strand, for the two converging forks at the RFB. Figure 5E shows that the 5’ end of the lagging strands of both converging forks stop some ∼50nt from RFB1. Importantly, the 3’ end of both leading strands proceed through the RFB and stop directly adjacent to the lagging strand of the opposing fork (Figure 5E). Thus, leading strands of the two converging forks appear to pass one another and are extended to meet the 5’ ends of the lagging strands ahead of them. Termination at RFB is likely a multi-step process where the rightward moving fork is first arrested and waits for the leftward moving fork to displace Fob1 – which is a polar barrier (Brewer et al., 1992; Hizume and Araki, 2019; Kobayashi et al., 1992). Removal of Fob1 by the leftward moving fork would permit CMG progression, allowing both leading strands to be extended until they reach the nascent lagging strands ahead of them. Our high-resolution analysis provides the first in vivo evidence to show that leading strands pass one another during the termination process (Dewar et al., 2015). It remains unclear why Rrm3 is needed for termination; at RFB, Rrm3 may promote Fob1 removal (Mohanty et al., 2006), but our data does not show distinct pausing of forks at RFB in *rrm3*Δ mutants. Figure 5E does show that the nascent ends of the converging forks are in close proximity to each other, thus it seems likely that small un-replicated gaps remain in *rrm3*Δ mutants (Deegan et al., 2019).

**Figure 5.**
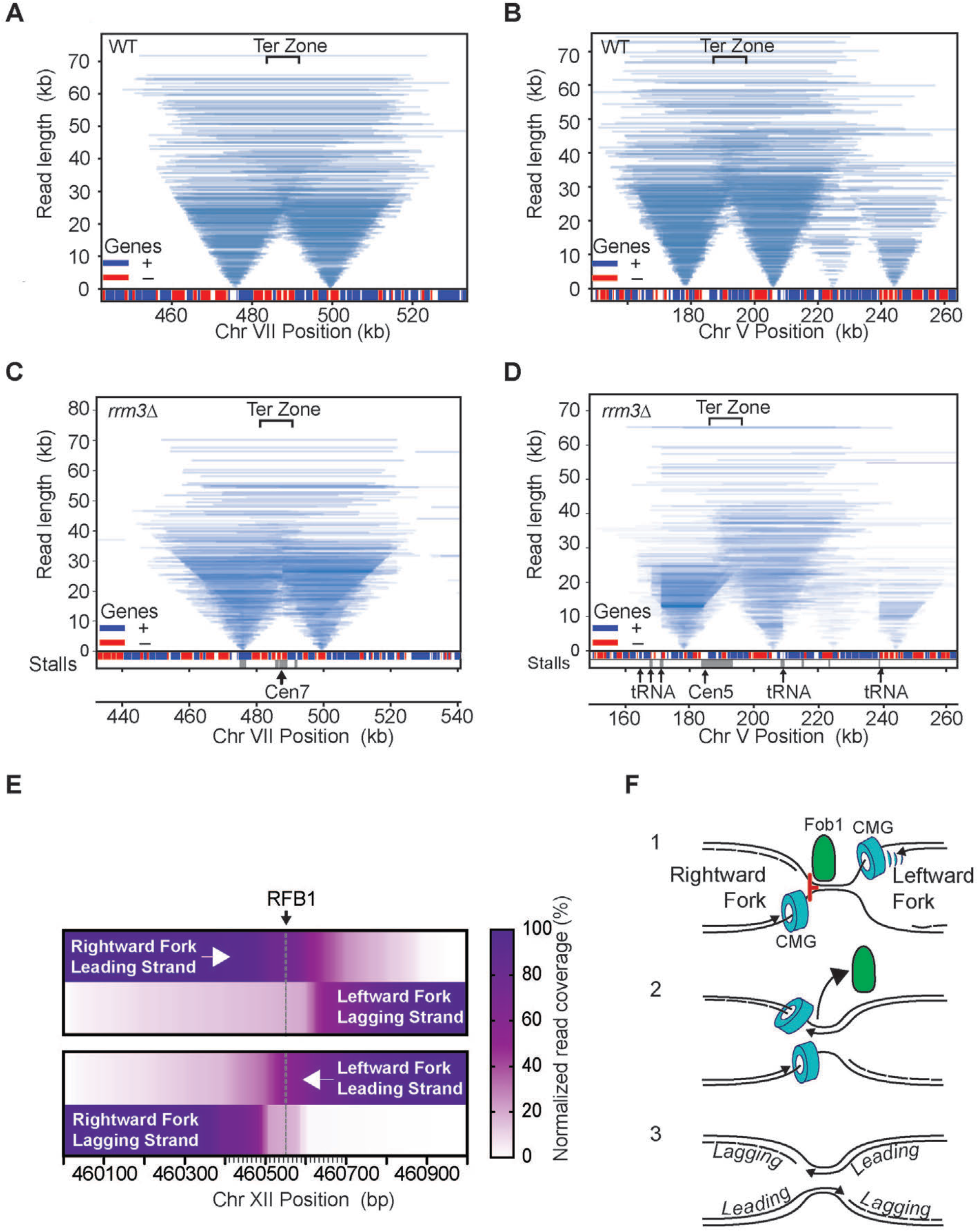
Tornado plot of replication termination zones. **A, B,** data from WT, **C** and **D** from *rrm3*Δ mutants. For clarity, only reads overlapping the replication origins are shown. Gene positions (red=Bottom, blue=Top strand) and stall sites in *rrm3*Δ (grey) are shown at the bottom. Prominent features are marked by arrows with text. **E,** Heatmap of read coverage of converging replication forks. Leading and lagging strands for left and right-ward moving forks are shown relative to RFB1. **F**, Model for replication termination at RFB site at the rDNA locus, see main text for details.

## Conclusion

The use of controlled nuclease digestion coupled with Nanopore sequencing provides a novel method for the analysis of coincident events on DNA. We have mapped the positions of sister replisomes in the process of DNA replication which has provided important new information on the relative rates of replisome progression the sites of replisome stalls, and the mechanics of replication termination.

We find that sister replisomes move through many kilobases of chromatin with surprising consistency, allowing us to calculate that the leading strand is typically ∼350nt longer than the lagging strand at advancing replisomes. This length is presumably dictated by size and rate of synthesis of Okazaki fragments, prior to their ligation into the nascent lagging strand. The high concordance of sister replisome progression also allows us to show that initiation of DNA synthesis occurs directly at the origin. Importantly, in both WT and *rrm3*Δ cells, arrest of one replisome is does not result in the arrest of the sister, demonstrating that the progression of sister replisomes is unlikely to be coordinated. Thus, despite their apparent colocalization (Kitamura et al., 2006), sister replisomes move independently but at a highly similar rate through a varying chromatin landscape. This finding illustrates that ancillary replisome factors, such as helicases and ATPases that promote fork progression through chromatin (Muellner and Schmidt, 2020; Shyian and Shore, 2021), are highly effective at mitigating the effect of potential impediments, including the transcription machinery (Gomez-Gonzalez and Aguilera, 2019). We confirm that Rrm3 is primarily required to overcome protein barriers, particularly TFIIIB at tRNA genes, centromeric nucleosomes and dormant origins(Azvolinsky et al., 2009), but we also find that Rrm3 is required for the replisome to overcome impediments in the promoter region of the *PDC1* gene. Presumably, this indicates that the RNA polymerase II pre-initiation complex is, under certain conditions, an obstacle to the replisome. Nevertheless, we found that S-phase transcription rate is not a strong determinant of replisome stalling in WT or *rrm3*Δ cells.

When stalling at RFB, we find that the leading strand advances to within ∼40nt of the impediment; while the precision of this measurement is lessened by our inability to define the exact position of the impediment, our measurements are consistent with previous reports and the finding that CMG prevents the leading strand from advancing to the fork junction (Duxin et al., 2014). Interestingly, despite the fact that the lagging strand template is not occluded by the helicase, the 5’ end of the lagging strand does not advance beyond the leading strand – indicating that unknown constraints prevent productive DNA synthesis on the lagging strand within ∼50nt of the replication fork. We find a similar pattern of nascent DNA end accumulation at tRNA and centromere stalls in *rrm3*Δ cells; however, in these cases, DNA end positions are far more heterogeneous than at RFB. This pattern may indicate that replisomes significantly slow or stall in the ∼200nt leading up to the DNA bound obstacle; or that a certain fraction of nascent DNA ends are processed following the prolonged stall.

Finally, we demonstrate that replicon-sequencing provides an unprecedented means to assay the process of replication termination. Our analysis shows that encroaching replisomes do not significantly slow as they approach each other, and that the sealing of converging replicons likely occurs without delay. Intriguingly, loss of Rrm3 activity results in a significant defect in replication termination at many sites across the genome. In part, this is explained by protein barriers acting as impediments to replisomes in termination zones (Fachinetti et al., 2010), however, even in the absence of protein barriers, converging replicons fail to efficiently join in *rrm3*Δ cells (Deegan et al., 2019). Such termination failure, though eventually resolved, results in a prolonged stall of the converging replisomes. The nature of the defect in *rrm3*Δ cells is presently unclear, but our data is consistent with biochemical data showing that Rrm3 and related helicases may be required to replicate through a small region separating two converging replisomes (Deegan et al., 2019).

Direct sequencing of DNA by Nanopore provides strand-specific information at nucleotide resolution, we anticipate that the roles of many replisome associated factors, key DNA processing events as well as chromatin assembly on nascent DNA can also be studied using this method.

## Limitations of the study

A primary limitation of this work is the relatively low throughput of the sequencing approach. While each sequencing experiment currently generates a ∼2-4 million sequencing reads, approximately 10% of these will be identified as BrdU containing and of use for analysis. Such numbers of BrdU containing reads generate high genome coverage, but replisome stalls and progression are best measured by quantitation of ends of nascent molecules. Thus, with the present throughput, we estimate that we can detect consistent stalls lasting more than 1 minute. Further optimization of the method should improve throughput and allow the detection of very infrequent or transient events. A second limitation of the study is our inability to precisely define how MNase cleavage influences the ends of nascent DNA molecules. While our data is consistent with previous studies that map ends of nascent chains, it remains possible that MNase cleavage is influencing the readout. Finally, we have focused on the analysis of DNA replication in early S-phase; as a result, our study is not designed to generate a comprehensive list of all possible replisome impediments across the yeast genome.

## Star methods

### Mnase-tag of MCM4

C-terminal fusion of codon optimized MNase was performed by transformation using pCC17 (MNase, yeast codon optimized version of pGZ108); MNase is separated from MCM4 by an 8 amino acid linker. N-terminal fusion of codon optimized MNase was generated using CRISPR technology. Guide RNAs for CRISPR and donor sequence are listed in Table S1

### Growth conditions

All strains used in this study are listed in Table S2. Yeast were grown in YPD at 25°C to OD_600_=0.35. Cells were arrested in G1 by addition of α-factor for 2h 40 minutes. Alpha factor was added each hour to a final concentration of 5ug/mL, 5ug/mL and 1ug/mL, respectively. 30min before the end of the arrest, the culture was supplemented with BrdU to 400ug/mL. Cells were washed 3 times with pre-warmed YPD and released in pre-warmed YPD supplemented with 400ug/mL of BrdU for 30 min at 25°C. 100mL of culture was pelleted at 4000rpm for 1 min, washed with ice-cold 20mM EDTA/1mM EGTA solution, pelleted at 4000rpm for 1min and flash frozen in liquid nitrogen.

### Nascent RNA-sequencing

Cells were grown and prepared in the same conditions as the Replicon-seq assay except that BrdU was omitted and 4-thiouracil was added to the media 15 minutes after alpha factor release for 15 minutes. Nascent RNA was extracted as described in the protocol (Baptista and Devys, 2018) with the addition of a mRNA selection step using NEB mRNA selection kit (NEB cat # E7490S) after streptavidin enrichment of nascent RNAs. Library was prepared using Nanopore direct C-DNA protocol (SQK-DSC109) and sequenced on MinION FLO-MIN106 flow cells R9. We performed two biological replicates and found a strong correlation between replicates (Figure S6C).

### MNase cleavage and library preparation

MNase cleavage protocol was adapted from (Grunberg and Zentner, 2017). Briefly, cells were thawed on ice with 1mL of ice-cold 20mM EDTA/1mM EGTA solution for 10min, pelleted for 20sec at 10,000g. Cells were washed 3 times with 1mL of Solution A. (For 5mL of solution A: 15mM Tris-HCl pH 7.5, 80mM KCl, 0.1mM EGTA, ½ Roche mini-PIC EGTA-free, 0.5mM spermidine and 0.2mM spermine). Cells were resuspended into 570uL of Solution A, incubated 1min at 30°C. 30uL of 2% of digitonin was added to the cells and incubated 5min at 30°C. Cell were incubated on ice for 2min, before addition of 2mM CaCl_2_. MNase digestion was performed on ice for 10 seconds before addition of 610uL of 2X stop buffer (400mM NaCl, 20mM EDTA and 8mM EGTA). Cells were pelleted for 20 seconds at 10,000g. Cell walls were removed with the addition of 5mg of Zymolyase 100T (Nacalai Tesque) in a 250uL of Zymolyase digestion buffer (1M sorbitol, 1mM EGTA, 10mM B-mercaptoethanol, and 50mM Tris-HCl pH 7.5) for 3min at 30°C. Spheroplasts were washed 2 times with Zymolyase buffer, and one time with RNase buffer (50mM NaCl, 10mM DTT, 50mM Tris-Hcl pH8.5). Resuspended in 500uL of RNase buffer and supplemented with 10uL of RNAse cocktail (Invitrogen), and incubated at 37°C for 2h. 2% final concentration of SDS, 50mg of chelex 100 resin (Biorad) and 10uL of 20mg/mL proteinase K (Goldbio) was added to the spheroplasts and incubated for 1h at 55°C. DNA was extracted 2 times by phenol-chlorophorm extraction in phase lock tubes, the phenol was gently mixed by inversion, and spun at 13,000rpm for 5min. The aqueous phase was transferred by pouring into a new clean tube. DNA was ethanol precipitated and resuspended overnight in 50uL of nuclease-free water.

Replicon-seq libraries were performed using SQK-LSK109 and SQK-LSK110 library kit from Nanoporetech. 1.2ug to 1.8ug of DNA of were used for library preparation. DNA repair and end-prep incubation time were changed to 10 minutes at 20°C. A-tailing was performed at room temperature for 15 minutes and elution from Ampure beads was performed at 37°C for 30min. 750ng to 1ug of library DNA was loaded on the MinION FLO-MIN106 flow cells R9, and sequenced for more than 48h, until pore exhaustion. We sequenced two biological replicates for WT and rrm3Δ mutant. WT run#1 we obtained 3.79.10^6^ basecalled reads, with a total of 1.03.10^5^ BrdU positive reads after BrdU calling by DNAscent, run #2 we obtained 4.87.10^6^ basecalled reads, with a total of 1.25.10^5^ BrdU positive reads after BrdU calling by DNAscent, rrm3Δ mutant run#1 we obtained 3.32.10^6^ basecalled reads, with a total of 1.39.10^5^ BrdU positive reads after BrdU calling by DNAscent, run #2 we obtained 4.31.10^6^ basecalled reads, with a total of 1.61.10^5^ BrdU positive reads after BrdU calling by DNAscent,

### Data analysis

All Fast5 files were converted into FastQ using Guppy (Oxford Nanopore Technologies). Sequencing reads were used to generate a custom reference genome using Canu (Koren et al., 2017), this allowed reliable mapping across transposable elements in the analyzed strain. All reads were mapped to the custom genome with Minimap2 (Li, 2018) using long read settings. BrdU containing reads were called using DNAscent v2 (Boemo, 2021). Using default settings, each read was split into 250bp intervals which were assigned as BrdU positive or negative. A BrdU score was then calculated for each read which represents the fraction of the BrdU positive intervals across the entire read. Only reads with a BrdU score ≥ 0.5 were used for analysis.

RNA seq data was converted to Fastq using Guppy and the mapped using Minimap2. The position and orientation of read primers were defined using LAST (Kielbasa et al., 2011) which allowed the orientation of the mRNA to be defined (Eccles, 2019). Median coverage for each gene was calculated using Bedtools (Quinlan and Hall, 2010).

### Genome coordinates, Data plotting

With the exception of the rDNA, all genome coordinates reported correspond the custom genome assembly. Liftover for genome coordinates was performed using RATT (Otto et al., 2011). Tornado plots were generated using custom python scripts available on request. Figure 4 C & D: Only singleton tRNA genes (not within 10kb of another tRNA) and reads overlapping annotated ACS sequences (Eaton et al., 2010) were used for analysis. Delta (Figure 4C) was calculated by first selecting reads with one end within 2kb of an annotated tRNA TSS. For each read, the distance from the tRNA proximal end to the closest overlapping ACS was calculated. The proximal distance was then subtracted from the distal distance (i.e. the distance from the distal read end to the ACS) to generate a value for delta.

### Stall sites and replication origins

Sites of replisome stalling were identified as regions with localized accumulation of read-ends. This was calculated by binning the genome into 200bp intervals and calculating the sum of ends for BrdU positive reads above 2kb in length in each interval. MACS2 (Zhang et al., 2008) was then used to call significant intervals using bdgpeakcall. Replication origins were identified by defining sites enriched for read midpoints. We mapped the midpoints of short reads (0-5kb) and long reads (5-20kb) to 200bp intervals along the genome. MACS2 was then used to call peaks using bdgpeakcall on short and long reads. We selected for peaks that overlap in both short reads and long reads as we found this reduces the bias introduced by the change in read midpoints during termination.

## Acknowledgements

We would like to thank Henry Cheng for work on analysis pipeline. M. Boemo, University of Cambridge, UK, for help with DNAscent. A. Viale and N. Mohibullah at the Sloan Kettering Integrated Genomics Operation for help with sequencing setup. D. Remus, X. Zhao, members of the Molecular Biology Program at SKI and members of the Whitehouse lab for comments and advice. **Funding:** This work was primarily supported by NIH grants R01RGM102253 and R01GM129058 awarded to IW. CC was also supported by an EMBO Long-term fellowship ALTF 246-2018. **Contributions:** Project was conceived by IW and CC, experiments were conducted by CC. Data was analyzed by IW, CC, JV and HC.

## Supplemetary material

### Supplementary figure legends

**Figure S1:**
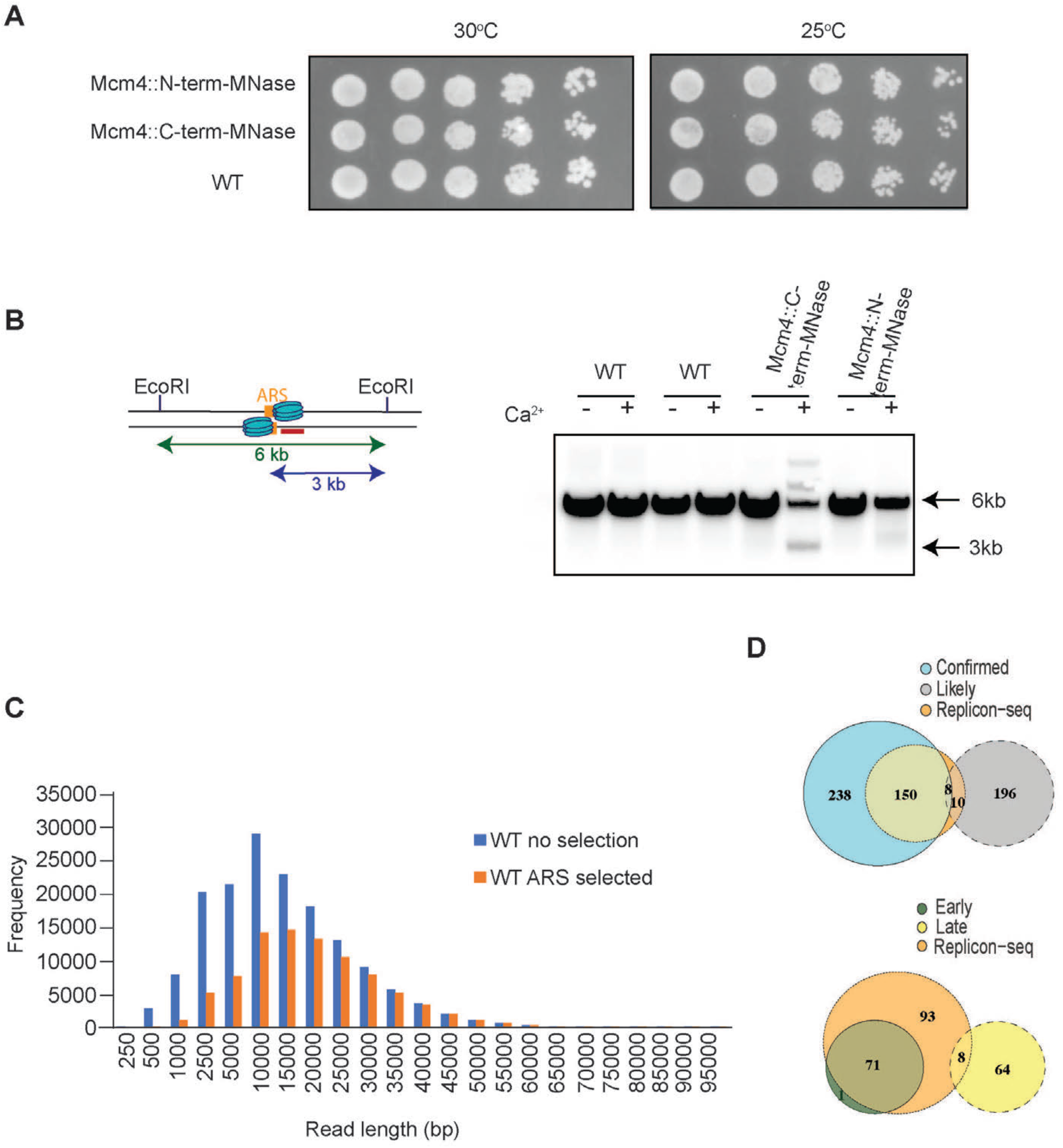
Mcm4-Mnase tag. **A**. Drop test showing no growth defect of MCM4-MNase fusions. **B**. The Mcm4-Mnase tag cleaves origin DNA in presence of calcium in G1. DNA was digested with *EcoRI* and probed with radiolabeled DNA binding to ARS305. In absence of cleavage by Mcm4-MNase 6kb band is observed, upon calcium addition Mcm4-MNase specifically cleave DNA at ARS releasing a 3kb band. **C,** Frequency plot of WT reads length, excluding rDNA. Blue bars show all BrdU containing reads, n=1.59 *10^^5^ (Methods). Orange bars show reads selected to overlap origins of replication (n=8.9.10^^4^). **D**, Venn diagram comparing Replicon-seq early S-phase replication origin calls (Methods) to known origins of replication. Total origins called: replicon-seq n=176, total OriDB (Confirmed) origins n= 401, late OriDB n=76, early OriDB n=72, likely n=214.

**Figure S2:**
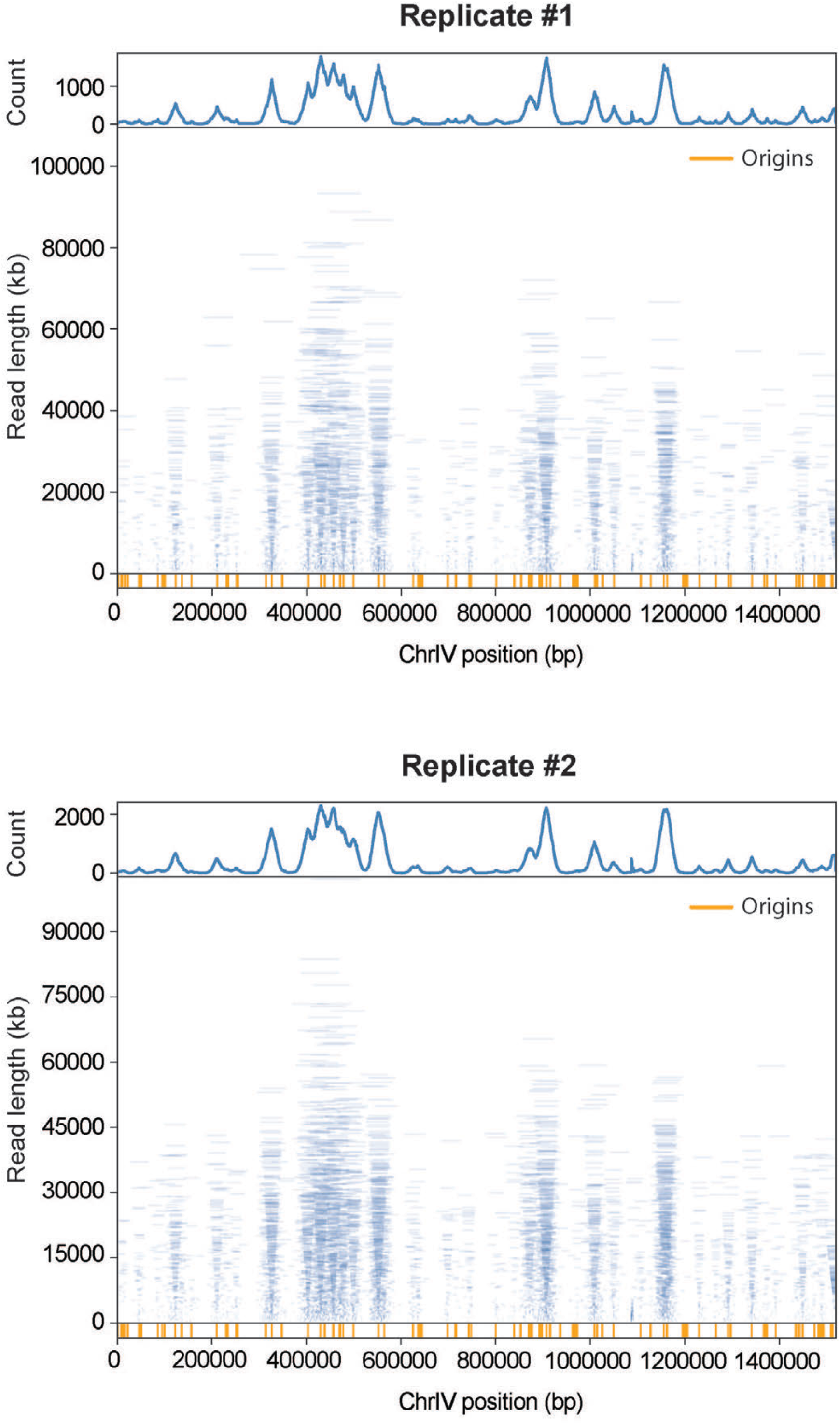
Tornado plots of WT Replicates.

**Figure S3:**
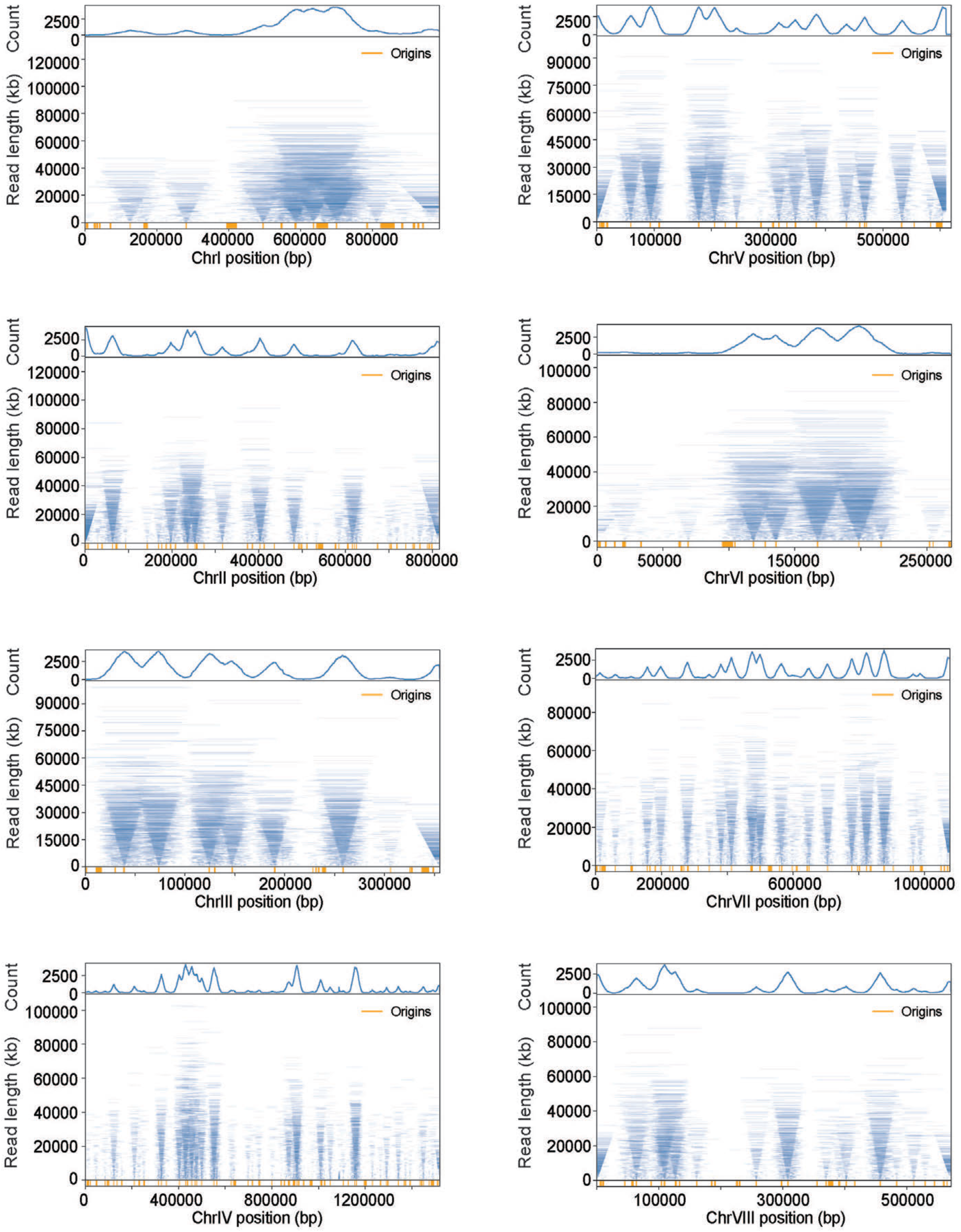
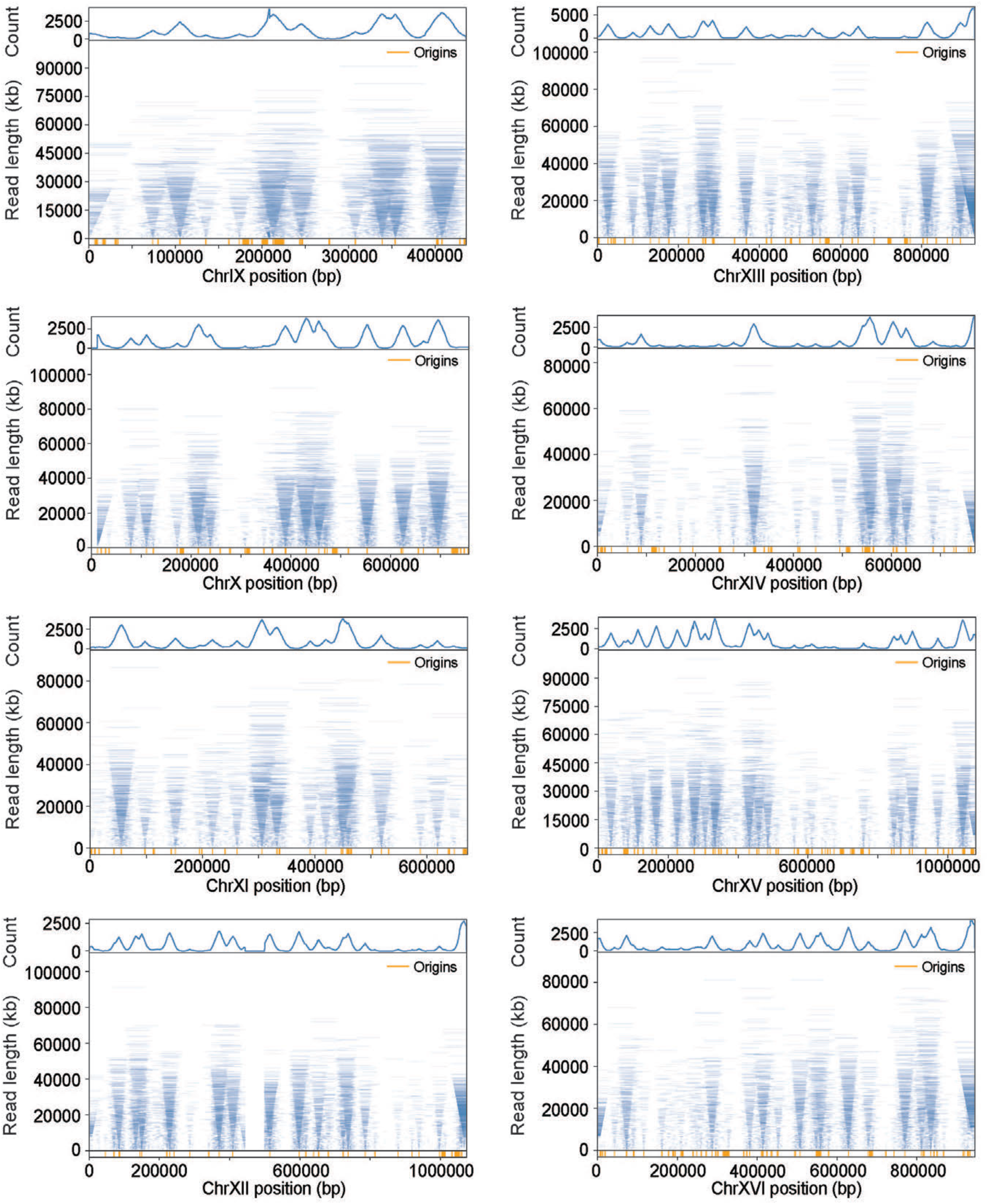
Tornado plots of all budding yeast chromosomes for WT cells.

**Figure S4:**
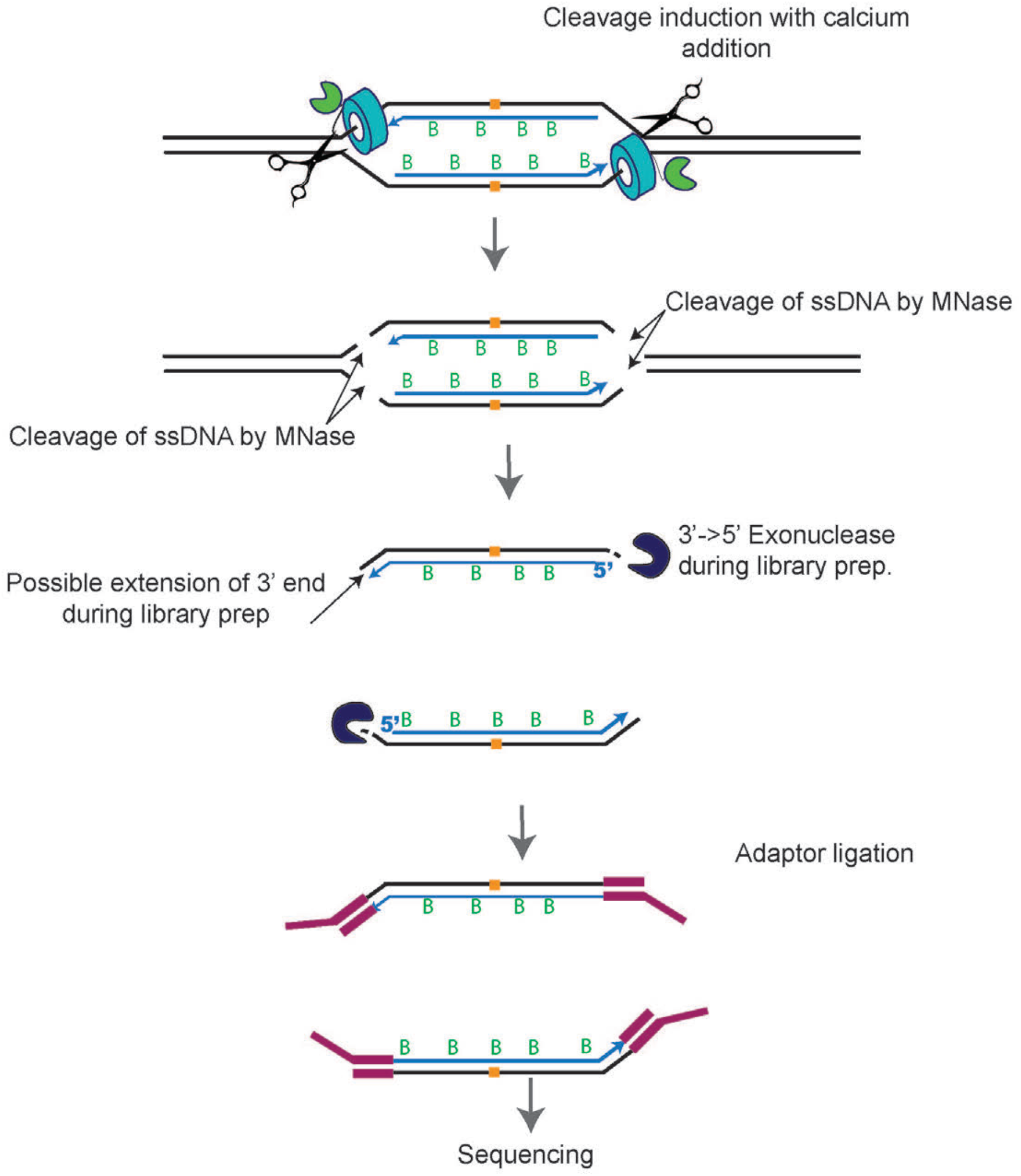
Schematic representation of MNase cleaving ssDNA at replication fork and library preparation for Replicon-seq. Cleavage of ssDNA at the replication fork should result in preferential digestion of the unwound parental DNA.

**Figure S5:**
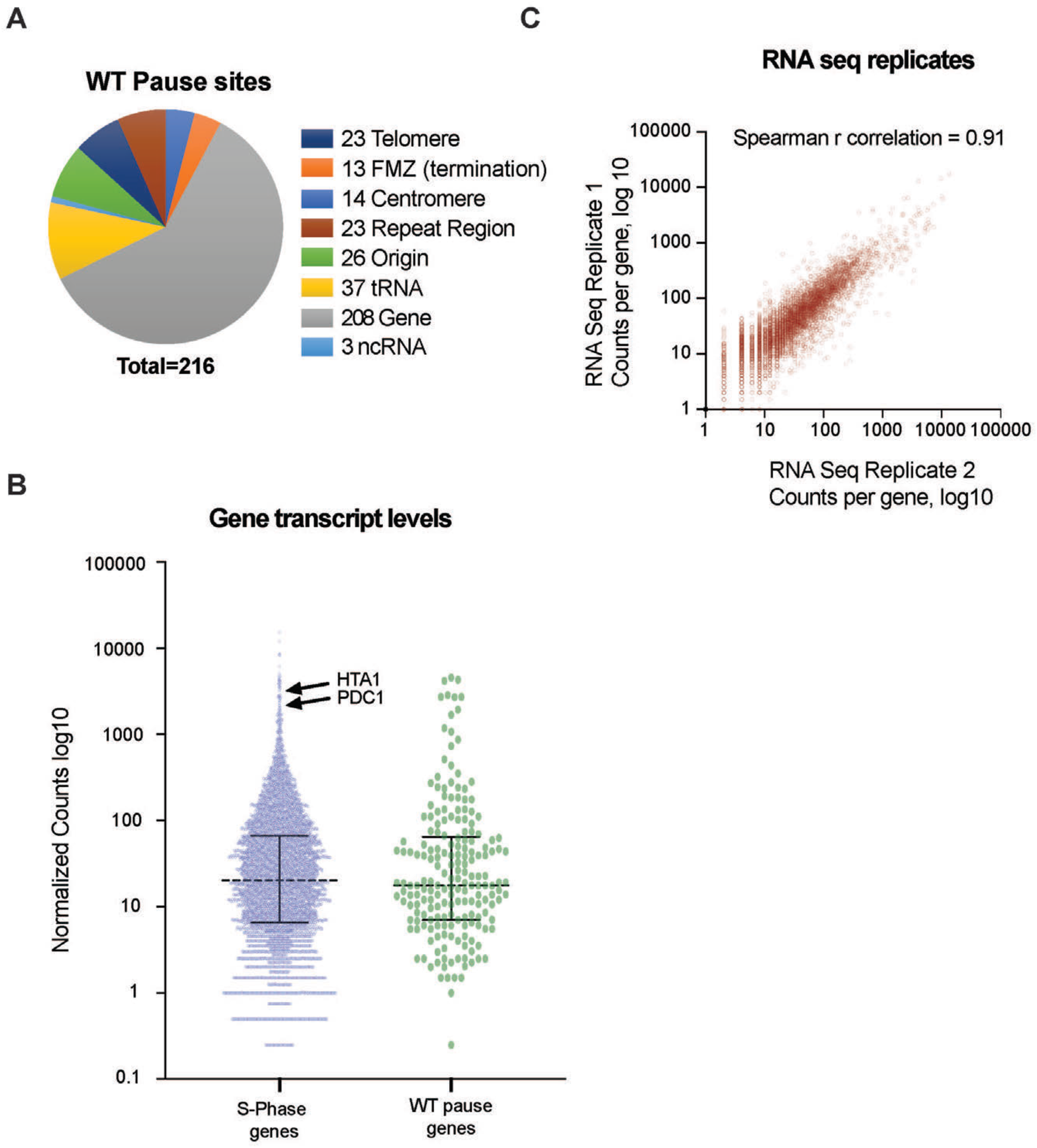
A, Chart showing the relative proportions of genomic features found at stall sites in WT cells. **B,** Scatter dot-plot of RNA seq transcription values for budding yeast genes. **C,** Replicates of Nascent RNA seq data for early S phase.

**Figure S6:**
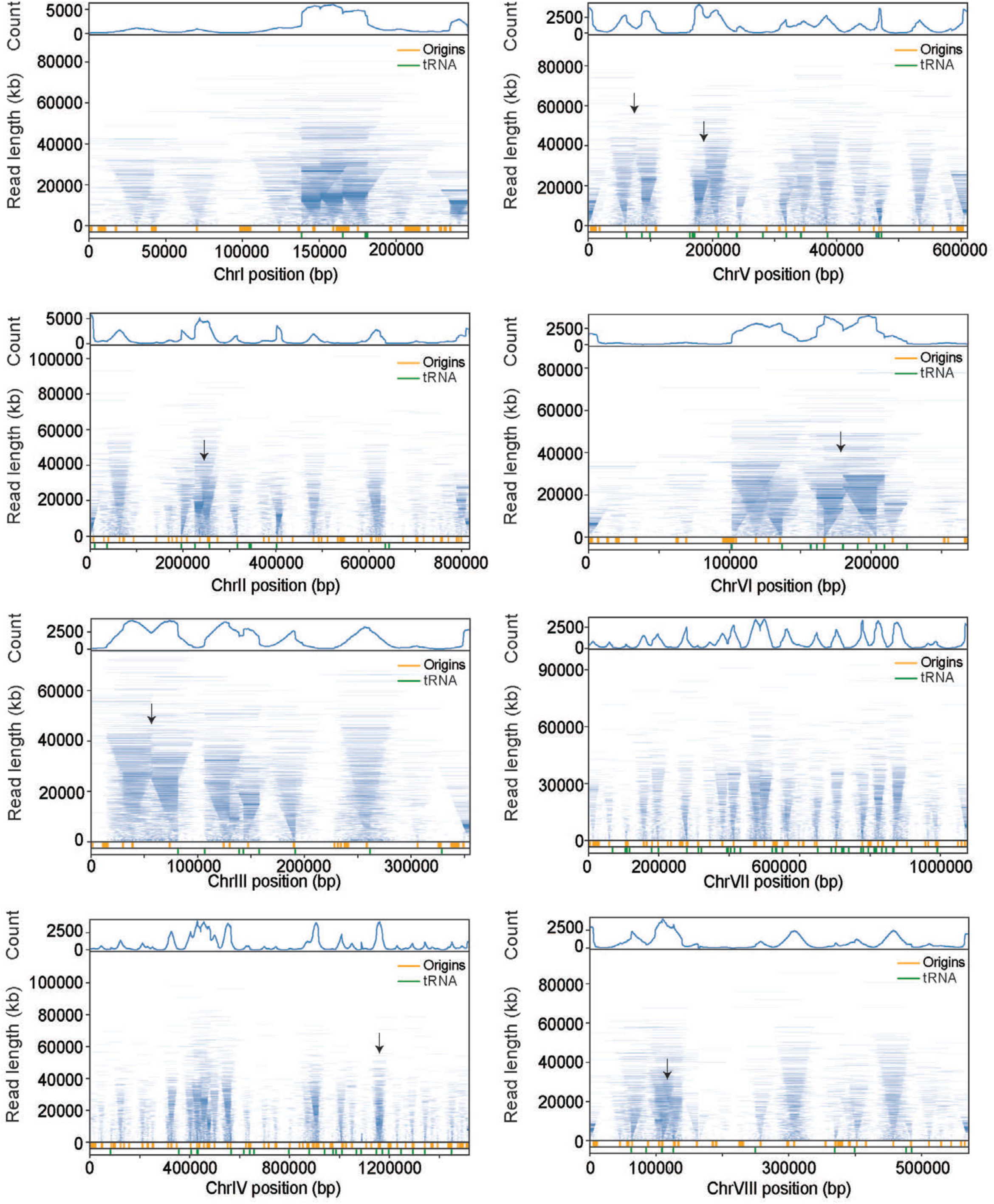
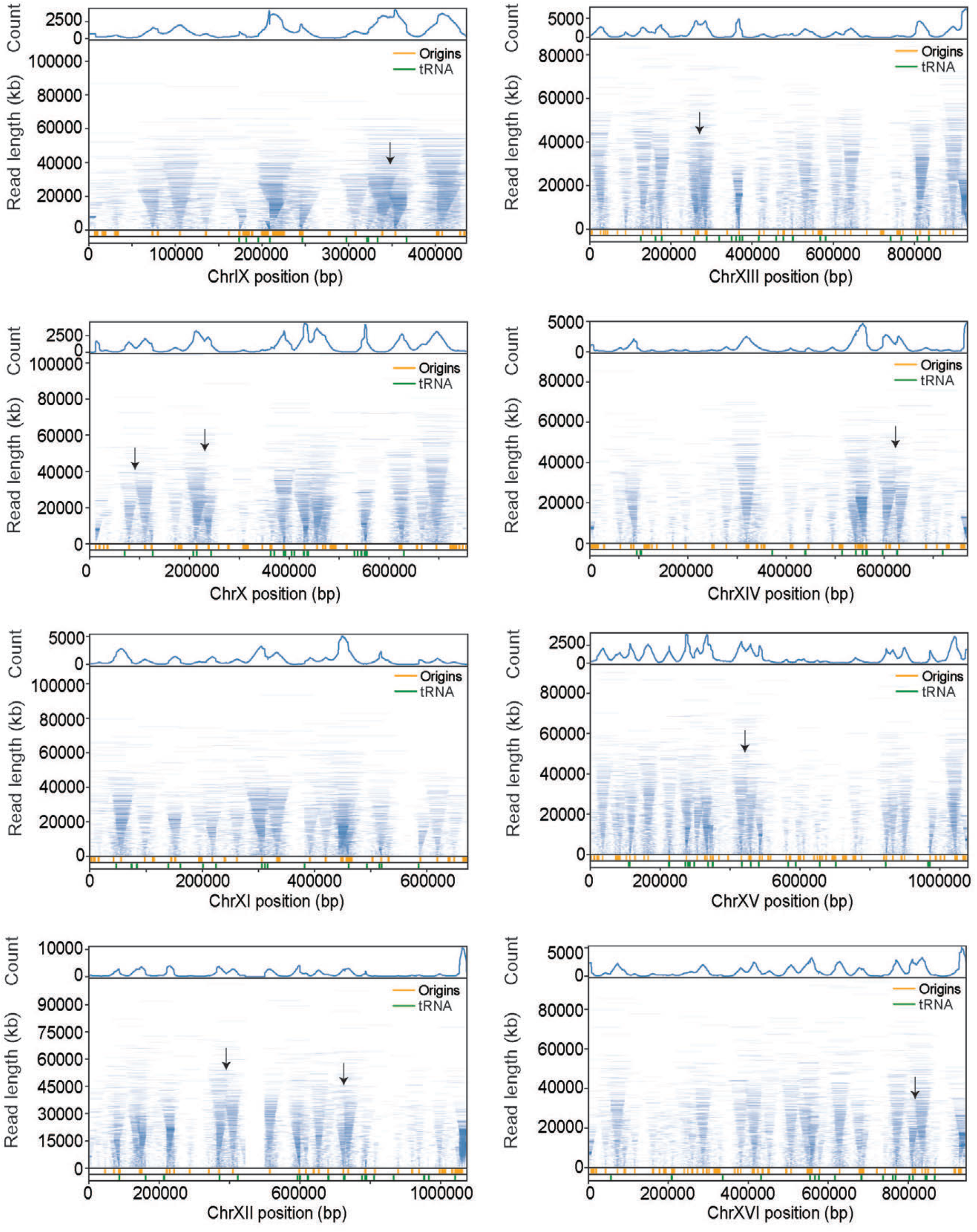
Tornado plots of all budding yeast chromosomes for *rrm3*Δ mutants. Black arrows indicate prominent sites of termination failure. Compare with Figure S3

**Figure S7:**
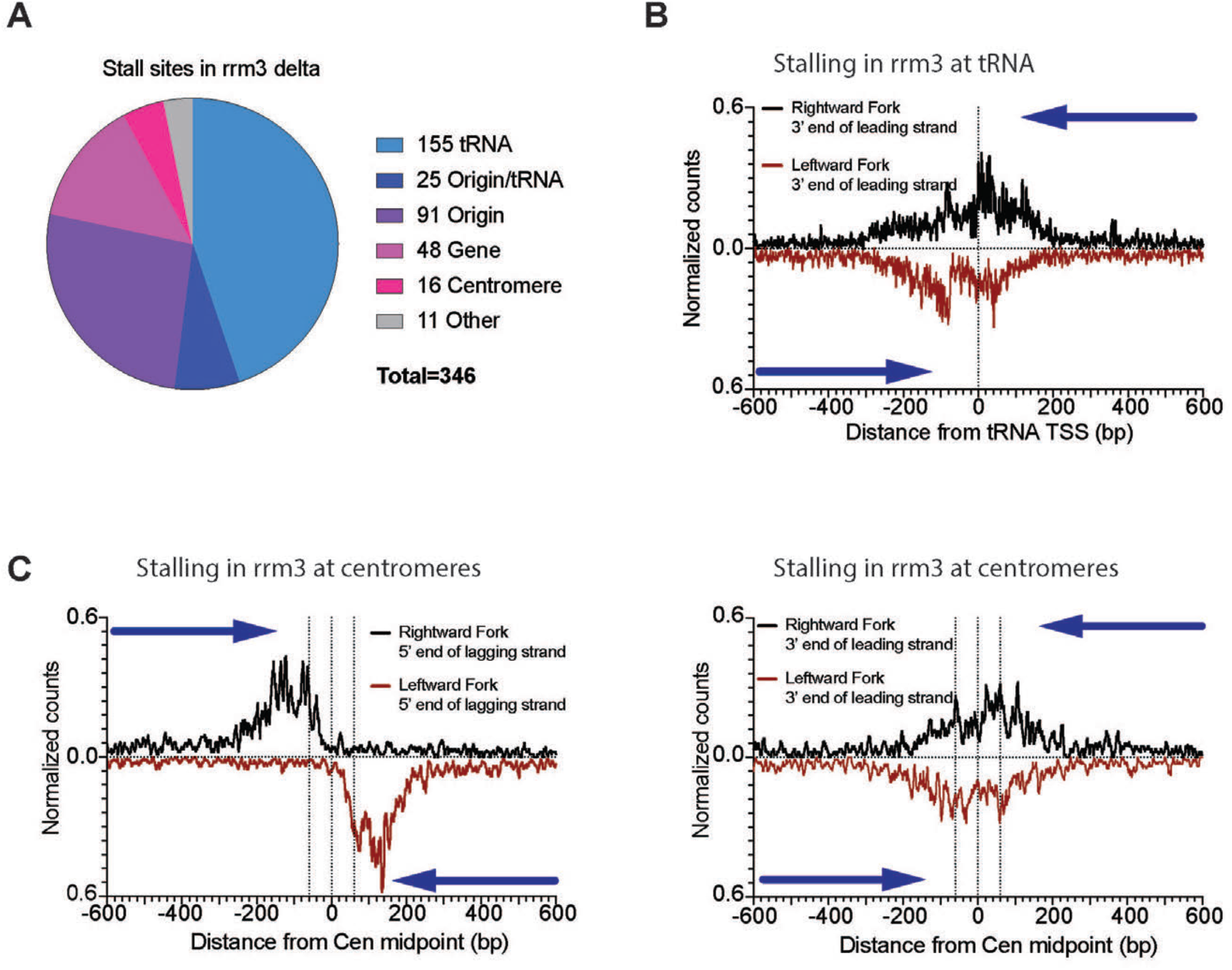
**A**. Diagram representing all the pausing site called from replicon-seq sequencing in *rrm3*Δ mutant in early S-phase. **B.** Meta-plot showing the distance of the 3’ of the rightward moving fork leading strand (black) or the leftward moving fork of the leading strand 5’ end (red) near the transcription site of 253 tRNA genes in *rrm3*Δ. Green dotted line shows the upstream edge of TFIIIB. **D.** Same as C, and Figure 4E for pausing at centromeres.

**Figure S8:**
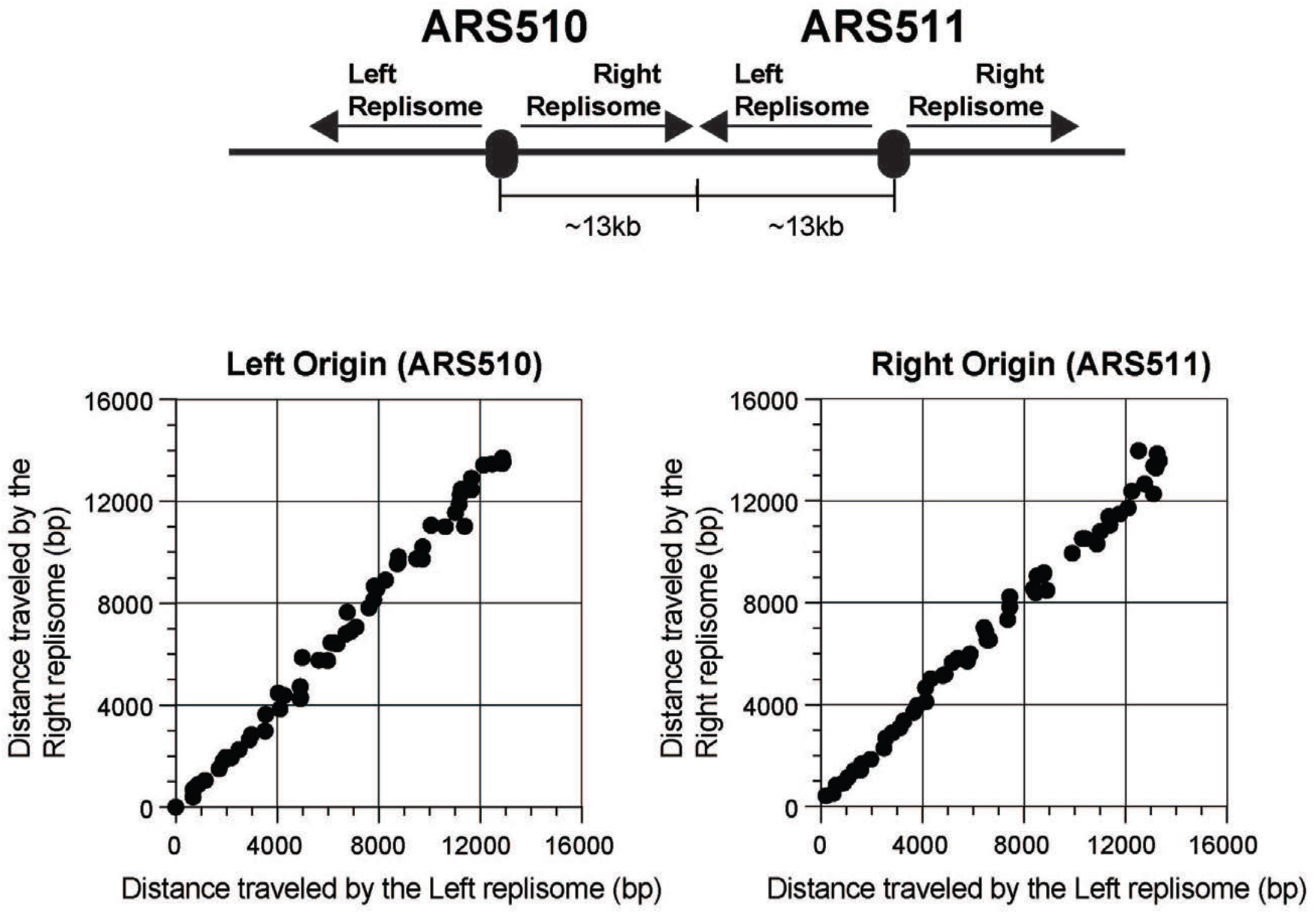
Relative progression of sister replisomes is plotted for Left origin (ARS510) and Right origin (ARS511). The Right fork of the ARS510 and the Left fork of ARS511 converge to terminate approximately 13kb from each origin. The median position of replisomes in 250bp intervals from the replication origin is calculated.

**Table 1:**
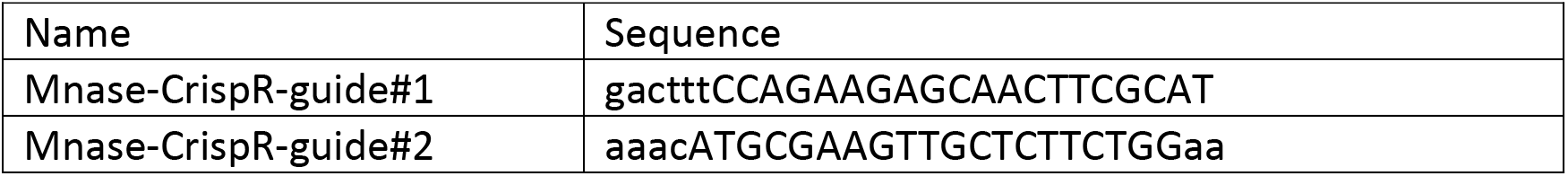
Guide RNA sequences used for the MCM4 N-terminal fusion by MNase.

**Table 2:** Strain genotype. All the strains are derived from W303 background (trp1-1, ade2-1 his3-11,15, can1-100).

| Strain name | Genotype |
| --- | --- |
| X7319 | MATa LEU2::BrdU-Inc URA3::GPD-TK7 |
| CCY285-6 | MATa MCM4-N-ter-MNase LEU2::BrdU-Inc URA3::GPD-TK7 |
| CCY295-1 | MATa MCM4-C-ter-MNase::KanMX LEU2::BrdU-Inc URA3::GPD-TK7 |
| CCY309-1 | MATa MCM4-N-ter-MNase rrm3ΔKanMX LEU2::BrdU-Inc URA3::GPD-TK7 |

**Sequence 1:**
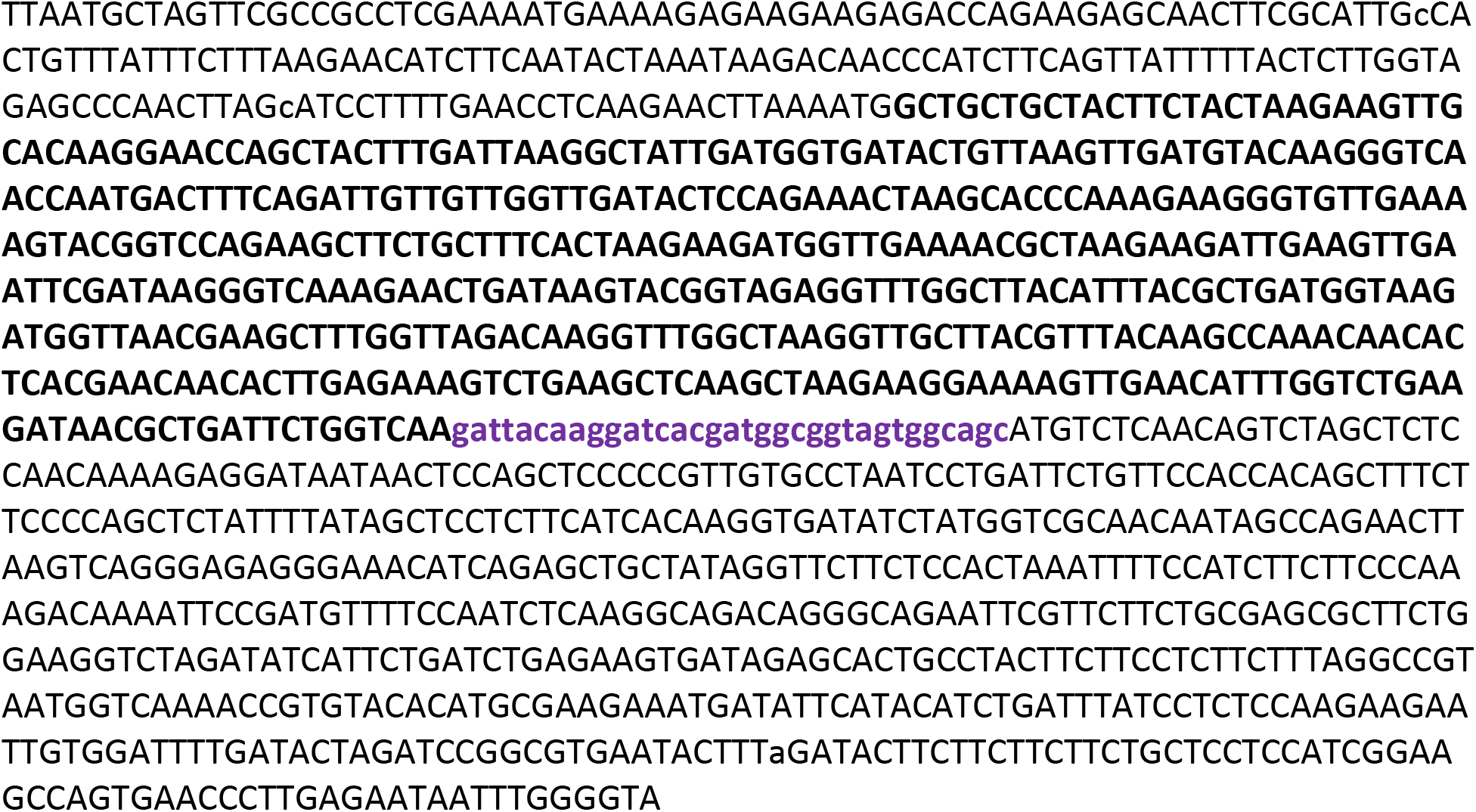
Sequence of yeast codon-optimized MNase used for the generation of the MCM4-MNase fusion protein.

